# Rooting density differentiates wheat genotypes through Bayesian modeling

**DOI:** 10.1101/089078

**Authors:** Anton P. Wasson, Grace S. Chiu, Alexander B. Zwart, Timothy R. Binns

## Abstract

Wheat pre-breeders use soil coring and core-break counts to phenotype root architecture traits, with data collected on rooting density for hundreds of genotypes in small increments of depth. The measured densities are both large datasets and highly variable even within the same genotype, hence, any rigorous, comprehensive statistical analysis of such complex field data would be technically challenging. Traditionally, most attributes of the field data are therefore discarded in favor of simple numerical summary descriptors which retain much of the high variability exhibited by the raw data. This poses practical challenges: although plant scientists have established that root traits do drive resource capture in crops, traits that are more randomly (rather than genetically) determined are difficult to breed for. In this paper we develop a Bayesian hierarchical nonlinear modeling approach that utilizes the complete field data for wheat genotypes to fit an *idealized* relative intensity function for the root distribution over depth. Our approach was used to determine *heritability*: how much of the variation between field samples was purely random versus being mechanistically driven by the plant genetics? Based on the genotypic intensity functions, the overall heritability estimate was 0.62 (95% Bayesian confidence interval was 0.52 to 0.71). Despite root count profiles that were statistically very noisy, our Bayesian analysis led to denoised profiles which exhibited rigorously discernible phenotypic traits. The profile-specific traits could be representative of a genotype and thus can be used as a quantitative tool to associate phenotypic traits with specific genotypes.

## 1. Introduction

Meeting the food production requirements of a growing human population who are encroaching on arable land and generating a changing climate will require an intensification of agriculture, where greater yields are obtained from crops on existing farms with sustainable inputs of water and fertilizer [1]. This will involve identifying the constraints on yield in agricultural systems; many of which are to be found below ground in the root systems of crops. There are calls for a “second Green Revolution” [2] focused on breeding crops with “ideotypic” [3] root systems (i.e. possessing desirable root system traits) that can overcome these constraints. This approach, called physiological breeding, is to be contrasted with breeding for increased yield alone, an approach which is no longer keeping pace with growing demand [4, 5, 6].

However, identifying ideotypic root systems for crops is fraught with difficulty. Root traits which can be identified in the laboratory are often difficult to translate to the field [7] because they are devoid of the developmental context of the soil. The soil environment is complex, and has a dominant effect on root system development [8]. It is also difficult to sample roots in soil in the field, and the results obtained are complex to interpret. Nevertheless, it is in the field where the effects of soil, climate, and agronomy are integrated with the developmental genetics of the plants, growing together as a crop, where measuring root traits and identifying crop ideotypes is most valuable. Selecting for root ideotypes in the field may speed up the identification of the best germplasm for breeding programs [8, 9].

Indirect measurements of crop root systems are problematic, and most direct measurements are destructive, time-consuming and/or labor intensive (e.g. root washing, minirhizotrons) [9]. Hence the core-break method was developed as a method of rapidly observing and quantifying the presence of roots as a function of depth [10, 11,]; a soil core sample is taken from the crop and broken at regular intervals (corresponding to increasing depth) and the exposed roots are counted. The counts correlate with the root length in the corresponding volume of soil. This technique has been used to phenotype root count distributions in 43 genotypes [12] and efforts have been made to automate the root count process [13] to reduce the labor requirements. However, root counts from the core-break method are subject to a high degree of variation between samples [11], which makes it challenging to identify genotype-specific traits from root counts or to associate genotypes with discernible properties of root count profiles.

Similar types of experimental field data may have been analyzed by statistical linear models [14] under an analysis-of-variance framework [12]. However, a major limitation of linear models is their assumption of Gaussian (normally distributed) response data, whereas root counts are discrete, bounded below by zero, and with a substantially right-skewed distribution even after any sensible variable transformation. Indeed, root count data are more adequately modeled as Poisson distributed, although a phenomenon known as overdispersion [15], commonly encountered in count data from field experiments, must be handled with care. More specifically, the Poisson distribution is characterized by a single parameter that represents the distribution mean as well as its variance. However, in practice, the count variable of concern often has a recognizably larger variance than its mean (hence, “overdispersion”), although the overall distribution still resembles Poisson in other respects.

Therefore, linear models applied to field data thus far have focused on analyzing core-level summary metrics, such as maximum rooting depth, which, after variable transformation if necessary, can approximately behave as Gaussian [12]. However, such summary metrics by definition cannot reflect root structure over depth, discarding valuable information contained at the level of individual core segments, and consequently resulting in an undesirable loss of statistical power.

To better facilitate our scientific objective of associating genotypes with discernible properties of root count profiles, in this paper we scrutinize the many facets of the inherent variability of the root count profile produced based on a field trial (Figure 1) that involved twenty genotypes (*n*_*G*_ = 20), each generating four replicated soil cores (*n*_*C*_ = 4) extracted in situ from each of four replicated plots or blocks (*n*_*B*_ = 4). Growing in a plot, as they would in a farmers field, the plants root systems interact and respond to each other. Their development is driven by the exploration of cracks and pores [16], which are randomly distributed. Likewise, variation in soil chemistry and nutrients, which can be patchy and vary with depth, drives the branching of roots. In contrast, impenetrable material and compaction can inhibit growth. As each soil core only captures a comparatively small piece of variation due to the various sources, results found in adjacent replicated cores can differ substantially.

**Figure 1:**
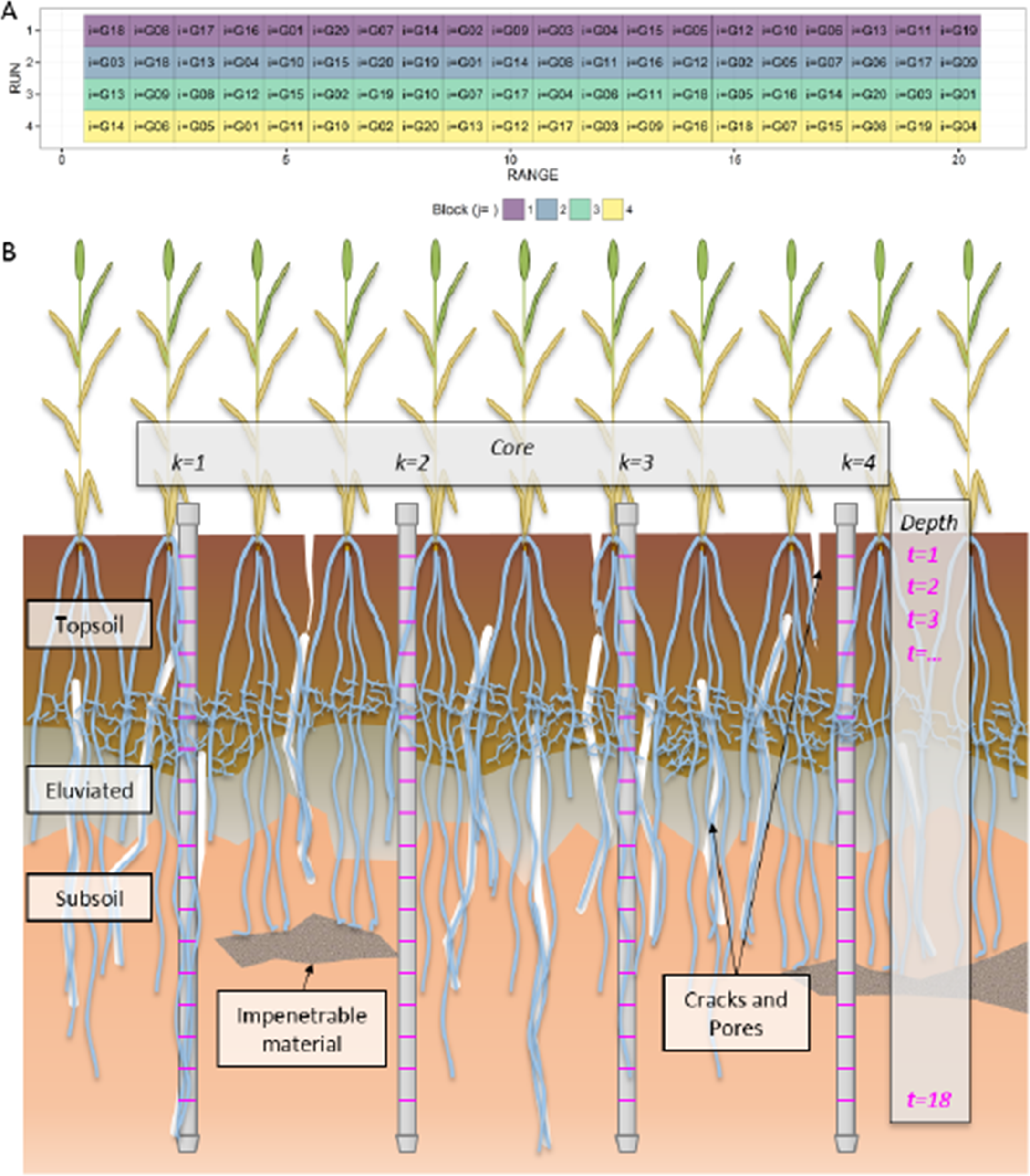
Schematics of the field experiment. **(A)** Surface layout of the field experiment involving twenty genotypes (indexed by *i*) randomized within four blocks (indexed by *j*) of twenty ranges of plots. **(B)** Cartoon depicting the sampling in each plot. Four soil cores (indexed by *k*) were sampled from each plot in a steel tube. Each core was broken into 10 cm increments. The root count (*y*) at each 10 cm depth increment (indexed by *t*) is the sum of the counts on the lower face of the upper fragment and the upper face of the lower fragment. Thus, each root count, *y*_*ijkt*_, has four unique index values. The cartoon further depicts the variability that might be encountered by sampling soil cores in a single plot, e.g. contrast cores *k* = 1 and *k* = 4.

Moreover, we note that the profiles of the average root count depicted in Figure 6 of ref. [12] strongly resemble the density function of the gamma probability distribution. Based on this observation, in this paper we develop a statistical modeling approach that can rigorously handle the non-standard nature of our root count data. Specifically, root counts at the observed depths (denoted by *t*) within a core are formally related through a nonlinear parametric expression *θ*(*t*) to reflect the one-dimensional spatial nature of individual soil cores. The parametric expression (with a small number of unknown parameters) is the common denominator that unifies this spatial behavior among all cores. Obviously, an appropriate parametric structure imposed on the root counts within a core would lead to much greater statistical power when compared to, say, an oversimplified analysis-of-variance approach that regards depth as a mere design feature in a factorial experiment.

We also note that our field experimental setup was such that the randomness in our data exhibits a hierarchical structure [17] that comprises layers of mean and variance functions. These features of our data require hierarchical nonlinear mixed modeling (HNLMM), an approach which addresses our need to model overdispersed, Poisson distributed data [15] via a hierarchy of nonlinear mean functions and associated variance components due to the formulation of *θ*(*t*).

We employ Bayesian modeling [18], an intrinsically hierarchical paradigm, to accurately reflect the complex, non-standard experimental design here. It is also greatly flexible: for a mathematically sound model, even with arbitrarily complex nonlinear parameters and random quantities that follow arbitrary probability distributions, rigorous statistical inference is straightforward once the model is computationally implemented.

Therefore, our approach in this paper is distinguished from existing studies particularly because of (a) our scrutiny of the root count profiles themselves, rather than the relationship between the counts and the root length density, and (b) our Bayesian modeling approach that integrates all identified facets of variability among all observed root count profiles in a comprehensive and collective manner. Additionally, our modeling framework gives rise to new heritability metrics that describe spatial and overall root architectural traits, the latter at the overall genotypic level.

## 2. Materials and Methods

### 2.1. Data and modeling framework

Each soil core sampled was partitioned in the field into five-centimeter segments from which the number of roots, *y*, was determined every 10 cm up to 180 cm using a fluorescence imaging system [13]. Each value of *y* at Depth *t*(= 1, …, *n*_*D*_ where *n*_*D*_ = 18) is the sum of the count imaged from the bottom face of the segment above *t* and that from the top face of the segment below *t*. (See Section 2.4 for details of data collection.) Let *y*_*ijkt*_ denote the total number of imaged live roots of Core *k* at Depth *t* for Genotype *i* in Block *j*. Thus, each ith genotype is associated with 288 (= *n*_*B*_*n*_*C*_*n*_*D*_) observations of *y* in total. Equivalently, each tth depth is associated with 320 (= *n*_*B*_*n*_*C*_*n*_*G*_) observed counts.

Data visualizations for Genotype G18 (Figure 2) and other genotypes (not shown) suggest that our observed root counts, *y*, perceivably follow a smooth nonlinear trend over core depth, but subject to substantive noise from the effects of soil physical and chemical properties described above, plus sampling and handling errors. These sources of noise culminate in the profile plots (Figure 2, panels A and B) and associated boxplots (Figure 2, panel A) for *y*. Therefore a modeling framework comprising the following main model statements was developed to capture the complex noise structure around an idealized smooth trend:

**Figure 2:**
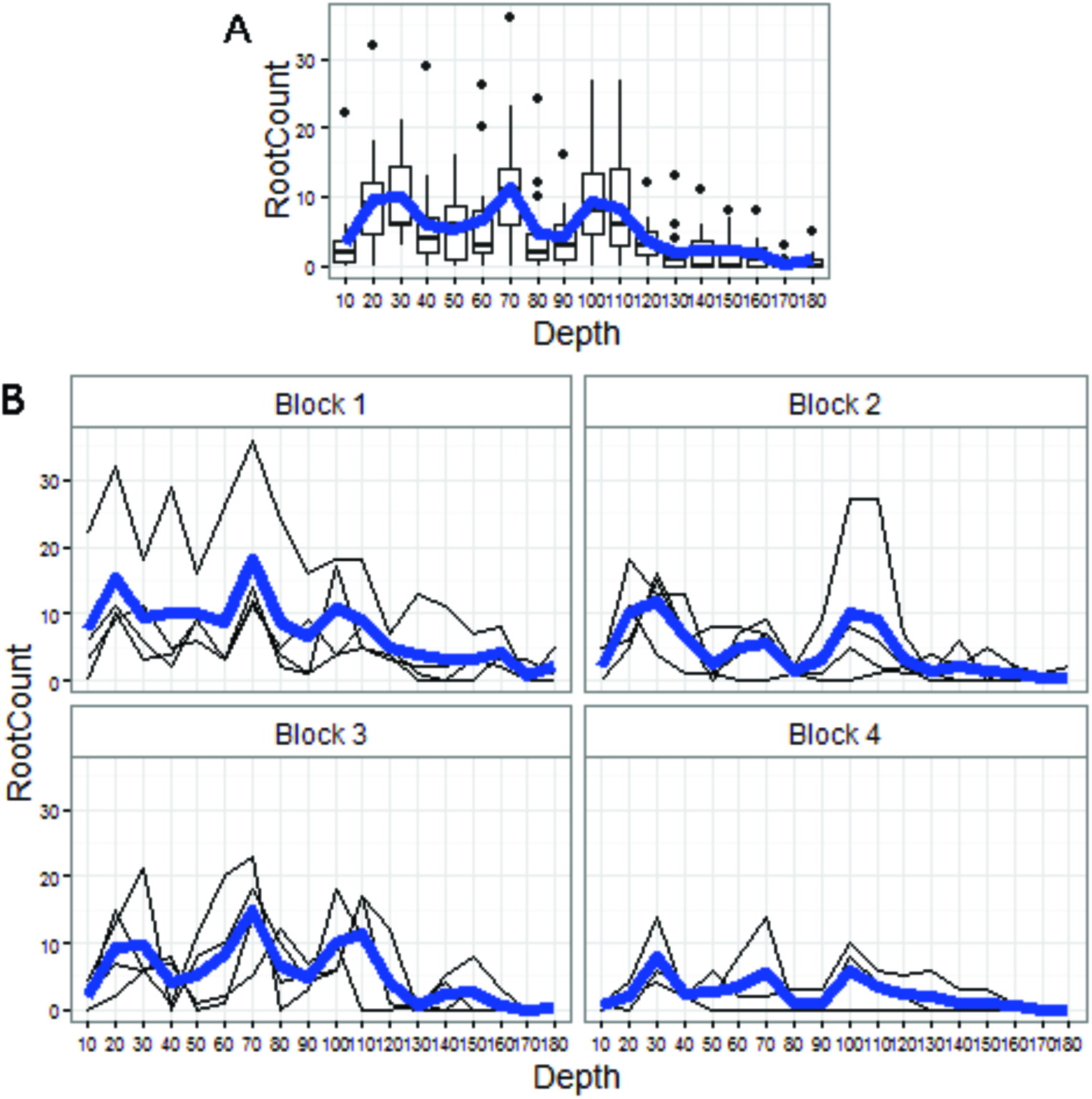
Data visualizations. **(A)** Boxplots of root counts, by depth for genotype G18, pooled across replicate plots (4) and depth-specific core segments (4 per plot). The horizontal axis is depth from 10 cm to 180 cm, at 10 cm intervals. The blue line is the empirical mean root count profile over depth, which, along with the corresponding mean profiles for other genotypes, strongly resembles those in Figure 6 of ref. [12]. **(B)** Root count profiles (in thin black) over depth, by block (replicate plot, shown as panel label), for genotype G18. Superimposed in bold blue within each block is the within-block empirical mean root count profile.

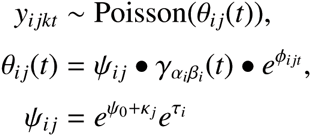

where *θ*_*ij*_(*t*) denotes the underlying plot-specific Poisson *intensity curve* over depth, i.e. the modeled mean root count at Depth *t* (= 1, 2,…, 18) from the {*i, j*}th plot (for Genotype *i* (= 1, 2,…, 20) observed in Block *j* (= 1, 2, 3, 4)).

Intensity *θ*_*ij*_(*t*) itself is modeled as being random, comprising a smooth genotype-specific “kernel function, *γ*_*α*_*i*_*β*_*i*__(*t*), and two sources of multiplicative Gaussian errors: genotype-specific deviation *τ*_*i*_ and core segment-specific deviation *φ*_*ijt*_. The block effects *κ*_*j*_ are modeled as fixed-effects parameters.

### 2.2 The root systems bulk and exploration parameters

The idealized function *γ*_*i*_(*t*) = *γ*_*α*_*i*_*β*_*i*__(*t*) = *t*^*α*_*i*_−1^*e*^-*β*_*i*_*t*^ has two genotype-specific parameters, *α*_i_ and *β*_*i*_, respectively representing the non-negative *shape* and *rate* of the gamma probability density function. Holding *β*_*i*_ constant and increasing *α*_*i*_ causes the ith kernel function to (a) peak at a lower depth and (b) exhibit more spread around the peak (Figure 3, panel A). Thus, *α*_*i*_ corresponds to both the depth at which the root system is most dense and its tendency to explore spatially around this depth. Henceforth, we refer to *α*_*i*_ as the *bulk parameter*.

**Figure 3:**
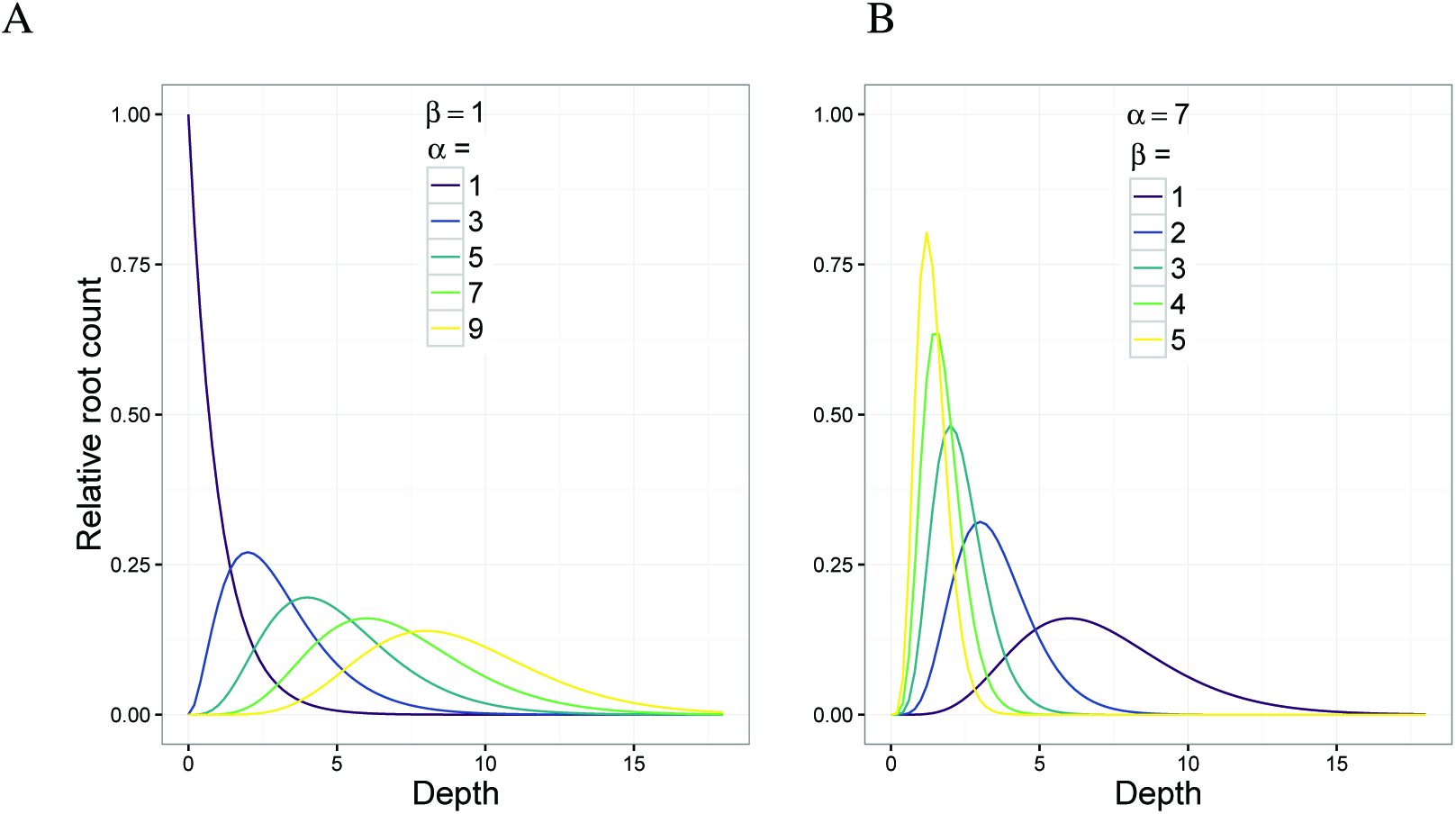
An illustration of the effect of changing the parameters *α*_*i*_ and *β*_*i*_ on the shape of the genotype-specific kernel function (*γ*_*α*_*i*_*β*_*i*__(*t*)) which is proportional to the probability density function of the gamma distribution. The vertical axis is the idealized relative root count (a dimensionless value). (**A**) Increasing *α*_*i*_ while fixing *β*_*i*_(= 1) causes the idealized functions peak to be located deeper under the soil surface and to be less concentrated. Thus, *α*_*i*_ is a *bulk parameter* that reflects the depth and density of the “bulk of the root system. (**B**) The effect of increasing *β*_*i*_ while fixing *α*_*i*_(= 7) causes an increased skewness in the tail of the idealized function, and consequently decreases the depth of the functions peak from the soil surface and increases its concentration. Hence, *β*_*i*_ is an inverse *exploration parameter*.

Similarly, holding *α*_*i*_ constant, increasing *β*_*i*_ causes the ith kernel functions tail to taper off more quickly, i.e. to exhibit a more slender tail (Figure 3, panel B). Thus, *β*_*i*_ roughly corresponds to the decline rate of the root systems downward exploration. In other words, the less slender (i.e. fatter) the kernel functions tail, the slower the decline of the root systems downward exploration (or, the bigger the tendency for the root system to explore downwards). Henceforth, we refer to *β*_*i*_ as the *exploration parameter*.

For each *i*th genotype, parameters *α*_*i*_ and *β*_*i*_ are modeled as bivariate log-normal random variables (i.e. they are bivariate Gaussian on the logarithmic scale with unknown correlation *ρ*). These parameters and the noise terms *τ*_*i*_ and *ϕ*_*ijt*_ are each modeled to have a mean that is constant across the study (i.e. not indexed by *i, j, k*, or *t*), and similarly for all the (co)variance parameters in the model (see Section 2.4).

The intensity functions proportionality multiplier *ψ*_*ij*_, on the logarithmic scale, represents the plot-specific intercept of the {*i, j*}th function. The intercept can be regarded as the modeled mean count (log scale) of the root system just below the soil surface. Therefore, *τ*_*i*_ corresponds to the genotypic random effect on this near-surface mean count. It is modeled as log-linear, where its mean can be expressed as a study-wide constant *ψ*_0_ plus a non-random block-specific shift *κ*_*j*_ (both taken to be fixed effects). (See Section 2.4.)

Finally, to complete the Bayesian inference framework, we specify reasonably non-informative prior distributions to reflect our lack of knowledge, in the absence of data, about the model parameters (see Section 2.4). Altogether, our Bayesian HNLMM as specified above is referred to as *Model 1*.

### 2.3. Novel heritability measures

The general notion of heritability is the proportion of phenotypic variation that can be attributed to genetics. Loosely, we have

> *P*henotype = *Genotype* + *Environment*,
>
> heritability = Var(*G*)/Var(*P*).

This definition of heritability assumes that genotypic and environmental variables are independent, linear components of the phenotypic response variable of interest. In practice, the biological notions of phenotype, genotype, and environment are abstract, and their quantifications that can be measured in an experiment may exhibit a complex co-dependence in a nonlinear fashion. Thus, a quantification of heritability that purely stems from a linear decomposition of the phenotypic response can be nonsensical in practical settings.

In the case that the measurable quantities and experimental design can be reasonably described using Poisson regression, ref. [19] adapts the linear (Gaussian) model-based definition of heritability to the scale of the linear predictor in a Poisson regression model, rather than the scale of the phenotypic response. Recently, this definition was extended to a temporal Poisson mixed model [20]. We further extend these ideas to define heritability measures based on segment-level count data.

Our adaptations below emphasize the challenge of detecting trends in root architecture from root count data that are both highly noisy and highly non-Gaussian, and that deviate substantially from a Poisson distribution; while considering data at a reasonably high spatial resolution may mitigate the challenge due to noise, it necessarily requires additional model complexity to address the non-standard statistical behavior of the data, and consequently, a novel quantification of heritability based on our new modeling paradigm.

As discussed in Section 2.4 below, the mean number of roots in *Model 1* is nonlinear in its parameters even on the logarithmic scale, and hence, it is not a linear predictor in the usual context of generalized linear models. Nevertheless, at each tth depth, we decompose the variability of log *θ*(*t*) into 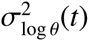, and 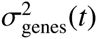 both of which are spatial in nature.

Here, we must address various aspects of complexity that are non-standard in heritability studies: (1) our analog of Var(*G*), namely, 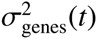, is attributable to the variability of the trio of genotypic parameters *τ*_*i*_,*α*_*i*_, and*β*_*i*_; and (2) it is a spatial function. Thus, it is reasonable to further decompose this Var(*G*) analog into *τ-*, *α-*, and *β*-specific components, as each of the trio pertains to different root architectural features; and the *α*- and *β*-specific components are also functions of *t* and are co-dependent except in the naïve case. In Section 2.4 below, we describe our four definitions of heritability (corresponding to 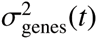, *τ*, *α*, and *β*) to handle such complexity.

Finally, we pool depth-specific values by taking the harmonic mean across depths, thus defining a quantity at the genotypic level that summarizes the particular architectural feature across all depths (see Section 2.4). The pooling of spatial elements to form an overall heritability measure gives rise to the multiresolution nature of our approach.

### 2.4. Technical details

#### 2.4.1. Data collection

The field trial was conducted at Ginninderra Experiment Station in Canberra, Australia (35°12′29.0”S 149°04′59.0”E), from late May to late December 2013 (the typical wheat growing season for the region) in alluvial cracking clay plots that were 1.3 m long. Twenty spring wheat genotypes (anonymized in this paper) were drawn from a collection of standard cultivars and from a multigenic mapping population on the basis of prior experimentation on root distributions in the field; each was sown with a tractor-drawn plot seeder in ten rows spaced 18 cm apart in a randomized block design with four replicated blocks of plots (Figure 1, panel A). A seed was sown roughly 3 cm apart in each row; the final sowing density was ~ 150 plants/m^2^. A fertilizer (N:P:K:S = 14:12.7:0.11) was applied at 120 kg/ha at sowing, with urea added for additional N during the growing season. Prophylactic fungicide and herbicide treatments were applied to the trial to suppress weeds and prevent disease. In early January 2014 after the trial had matured and been harvested, 4 soil cores of ∽1.8 m in length were collected from each plot using 2 m long, 42 mm diameter stainless steel coring tubes driven into the soil vertically with a tractor-mounted hydraulic push press [12]. Our field sampling technique ensured that within each plot the cores were reasonably independent of each other (Figure 1, panel B). Each core was broken into segments rather than sliced, so that the roots traversing the plane of the break would emerge intact from one of the two broken faces; the same root could not be visibly intact on both faces simultaneously. (Slicing the roots would have left only the cross sectional area on the face: 50–150 microns in diameter and difficult to detect.) Hence, the root counts on the adjoining faces can be regarded as independent values which, when combined to form y, represent the number of roots traversing the break plane at that depth. The fluorescence imaging system generates root counts [13] which necessarily differ from an observers manual counts, although both are subject to measurement error. The raw imaging data were processed (available from Supplementary Material) and visualizations produced with the statistical programming language **R** [21] using the packages ‘dplyr [22] and ‘ggplot2 [23].

#### 2.4.2. Formulation of Model 1 and variants

Recall that *y*_*ijkt*_ denotes an observed root count, where {*i, j, k, t*} indexes {genotype, block, core, depth}, fori ∊ {1,2,…,20}, *j* ∊ {*1,2,3,4*}, *k* ∊ {1,2,3,4}, and *t* ∊ {1,2,…, 18}. Taking *γ_i_(t)* = *t*^*α*_*i*_−1^ *e*^−*β*_*i*_*t*^ which is the kernel of the gamma probability density function and letting “N” and “BVN” respectively denote “normal” and “bivariate normal,” our model statements can be rewritten as follows:

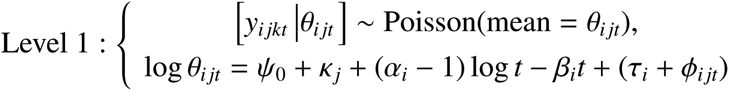

where

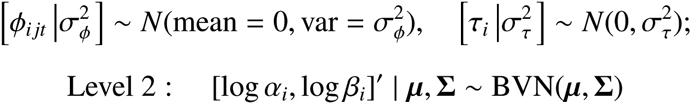

where

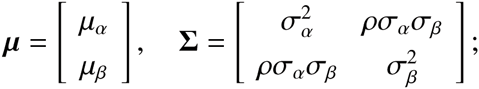

and *k*_*j*_*s* are fixed effects that require a linear constraint to ensure model identifiability: we take *k*_4_ = 0.

For Bayesian inference, prior distributions are required for all fixed-effects parameters *κ*_*j*_*s*, *ψ*_0_, *μ*_*α*_, *μ*_*β*_, and (co)variance parameters 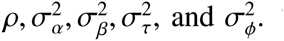 To reflect our lack of *a priori* insight (in the absence of data) into the likely values of these parameters, each was given a popular diffuse prior: the Fisher-transformation *arctanh*(*ρ*) and fixed effects were all assumed to be independent zero-mean Gaussians with a variance of 10^4^, and the variance parameters were assumed to follow independent and identical inverse-gamma(*a* = 1, *b* = 0.1) distributions. The resulting Markov chains of posterior draws exhibited very poor mixing for *Model 1* (as well as *Model 2* by prespecifying *ρ* = 0) when a smaller value of *b*, namely, 0.01 or 10^−4^, was used. As smaller values of *b* correspond to more diffuse inverse-gamma priors, the poor mixing suggests mildly weak identifiability (even for the smaller *Model 2*). This also suggests that to improve inferential power for and the identifiability *of Model 1*, one could conduct a future field study that consists of a larger number of plots and/or depths, and/or employ stronger priors based on the inference we have presented in this current article.

For model validation (discussed in Section 3), we also considered smaller models: *Model 2* by prespecifying *ρ* = 0 in *Model* 1; and *Model 3* by taking all *Model 2* parameters (including those that are genotype-specific) to be fixed effects.

#### 2.4.3. Multiresolution heritability under Model 1

To handle the nonlinearity ― even on the logarithmic scale ― of the mean number of roots *θ* in *Model 1*, we decompose the variability of log *θ*(*t*) into

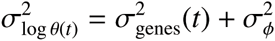

where

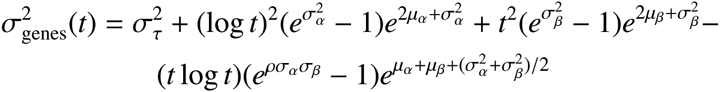

is attributable to the variability of the trio of genotypic parameters *τ*_*i*_, *α*_*i*_, and *β*_*i*_, while the study-wide parameter 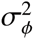 is attributable to the pure noise term *ϕ*_*ijt*_. Note that the parameters 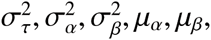 and *ρ* stipulate the collective statistical behavior, *a priori*, of *τ*, *α*, and *β*.

As such, we define four different measures of heritability, namely, 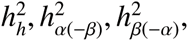 and 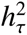 each at the genotypic level, by letting

> 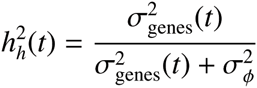 = depth-specific heritability of intensity function at *t*,
>
> 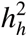 = heritability of overall architecture
>
> = harmonic mean of 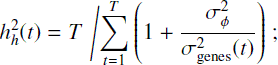
>
> 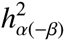 = heritability of root bulk’s location (and size) on log scale, IGNORING its relation with penetration rate
>
> = harmonic mean of 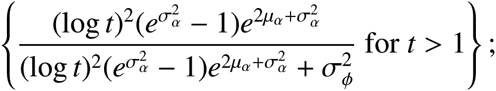
>
> 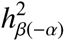 = heritability of root’s decline rate of penetration on log scale, IGNORING its relation with bulk location
>
> = harmonic mean of 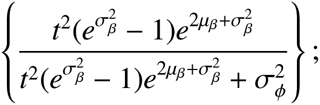
>
> 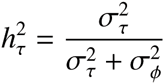 = heritability of intensity function’s intercept on log scale.

Note that each of 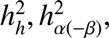, and 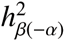comprises depth-specific heritability components, but 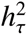. does not (and thus, its definition does not require the use of the harmonic mean).

#### 2.4.4. Implementation of Model 1 and variants

Bayesian inference requires the derivation of the joint posterior distribution of all model parameters. In our case, this distribution is intractable and Markov chain Monte Carlo (MCMC) was used to approximate it. For this, we used the RStan MCMC software [21, 24,] to fit *Model 1* and its variants for model refinement purposes (see Supplementary Material). Of the 320 cores sampled, nine were ignored as they failed to yield root count data. While missing data can be imputed under extra model assumptions, the Stan framework did not yet readily allow simulation of discrete parameters, and thus imputation of the missing root counts was not performed.

## 3. Results

### 3.1. Biological insights according to the statistical inference for Model 1

The following results are based on the joint *posterior distribution* among the parameters of *Model 1*.

#### 3.1.1. Root intensity profiles are statistically distinguishable among genotypes

Posterior inference allows us to examine the intensity profiles *θ*_*ij*_(*t*) and their idealized (de-noised) counterparts *ψ*_*ij*_*γ*_*i*_(*t*) for any given replicate block *j*. A *ψ*_*ij*_*γ*_*i*_(*t*) profile is effectively the intensity profile *θ*_*ij*_(*t*) but ignoring the random genotype-block interaction *φ*_*ijt*_. Note that *ψ*_*ij*_ is log-linear in *τ*_*i*_ and *κ*_*j*_ without an interaction term. Thus, the behavior among the 20 idealized profiles within any *j*th block is necessarily consistent across all four blocks but for an intercept shift *κ*_*j*_. Hence, Figure 4 focuses on *j* = 1 to represent the study-wide behavior of the idealized profiles.

**Figure 4:**
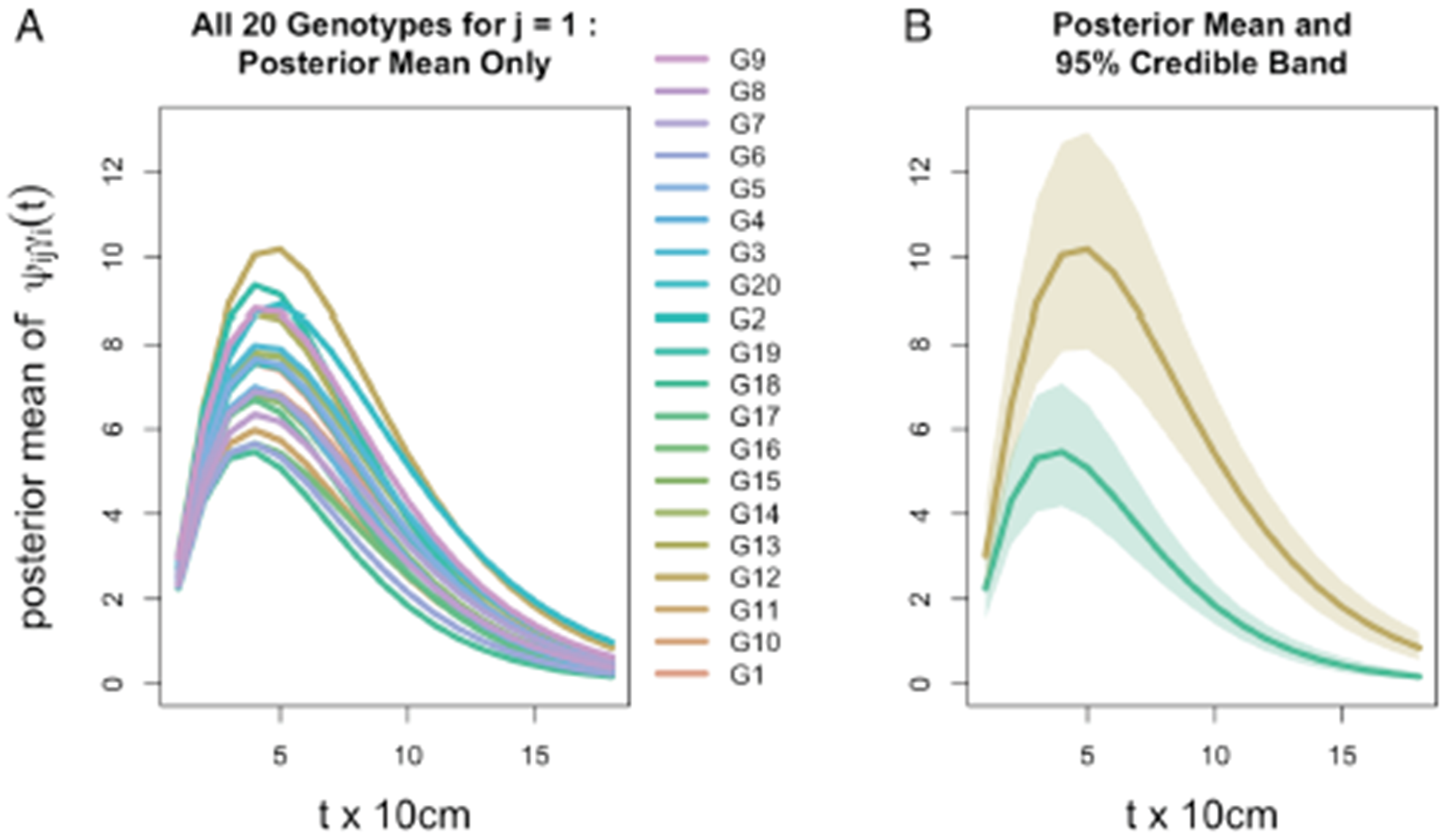
Inference for root distribution. (**A**) Posterior mean (Bayesian estimate) of idealized intensity profile *ψ γ*(*t*) for replicate block *j* = 1 for all 20 genotypes. Other blocks appear similarly, differing only in the intercept due to the block-specific fixed effect *κ* in which log*ψ* is linear. (**B**) Posterior means for Genotypes G12 and G18 (from panel A) which are respectively the maximum and minimum curves, each surrounded by a 95% credible band (Bayesian confidence band). Note that credible bands are constructed from depth-wise 95% credible intervals of *ψ*_*ij*_*γ*_*i*_(*t*); thus, the upper band limit is constructed by connecting, across the 18 values of *t*, the 2.5th percentiles of the *ψ*_*ij*_*γ*_*i*_(*t*) posterior distribution; similarly, the lower band is constructed by connecting the corresponding 97.5th posterior percentiles.

All posterior means (Bayesian estimates) of the 20 genotypic idealized profiles *ψ*_*ij*_*γ*_*i*_(*t*) are visually distinguishable (Figure 4, panel A); and the study-wide statistical power is very high in determining that the genotypes do not all exhibit the same idealized profile (Figure 4, panel B): 95% credible bands (Bayesian confidence bands) around the maximum and minimum idealized profiles are clearly non-overlapping. The lack of overlap at such a high credible level indicates that, among the 20 genotypes, at least G12 and G18 are highly statistically discernible with respect to their idealized intensity profiles.

While *ψ*_*ij*_*γ*_*i*_(*t*) necessarily behaves similarly across all *j*, the genotype-block interaction intrinsic in the plot-specific intensity profile *θ*_*ij*_(*t*) induces variability in the 20 profiles collective behavior across *j*, as is evident in Figure 5: in each block, this variability reduces the statistical distinguishability among the 20 genotypes, although in each of Blocks 2, 3, and 4, at least two intensity profiles are highly discernible. Specifically, despite the noisy nature of *θ*_*ij*_(*t*), Figure 5 shows that in each of Blocks *2–4*, at least two intensity profiles *θ*_*ij*_(*t*) (respectively, (*i* =){G2, G17} in Block (*j* =)2, {G6, G13} in Block 3, and {G6, G15} in Block 4) are highly statistically discernible due to the general lack of overlap between the pair of block-specific 95% credible bands around *θ*_*ij*_(*t*).

**Figure 5:**
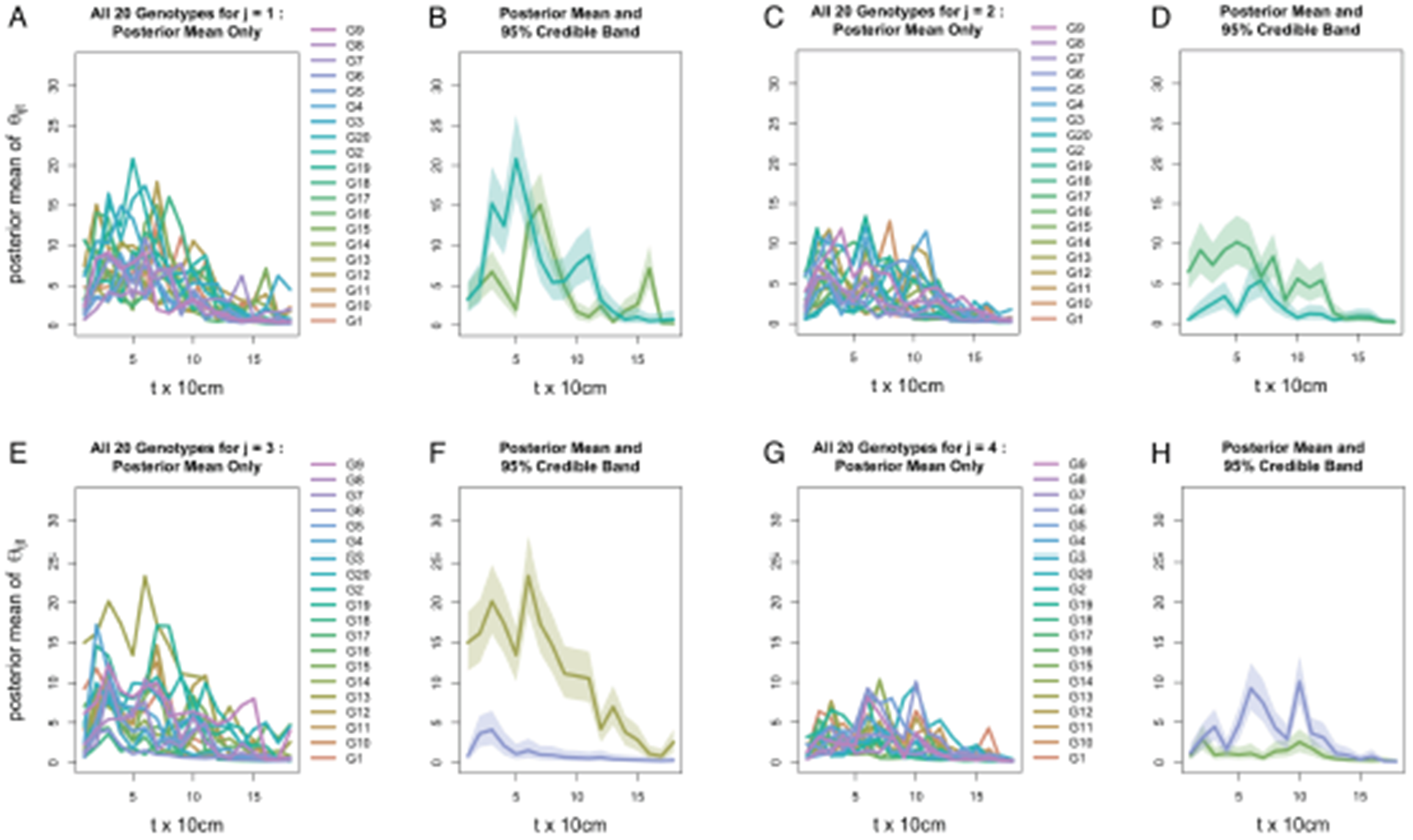
**Posterior mean of intensity profile *θ*(*t*) for all 20 genotypes**, coupled with those genotypes with the maximum and minimum curves and their 95% credible intervals for *j = 1 (**A, B**), *j* = 2 (**C, D**), *j* = 3 (**E, F**), and *j* = 4* (**G, H**).

#### 3.1.2. Root intensity profiles are substantially heritable

Each of our four genotypic heritability measures is a model parameter that exhibits a posterior distribution, shown in black in Figure 6; three of these are pooled measures, each comprising 18 depth-specific components (see Section 2.4 above), shown in color.

**Figure 6:**
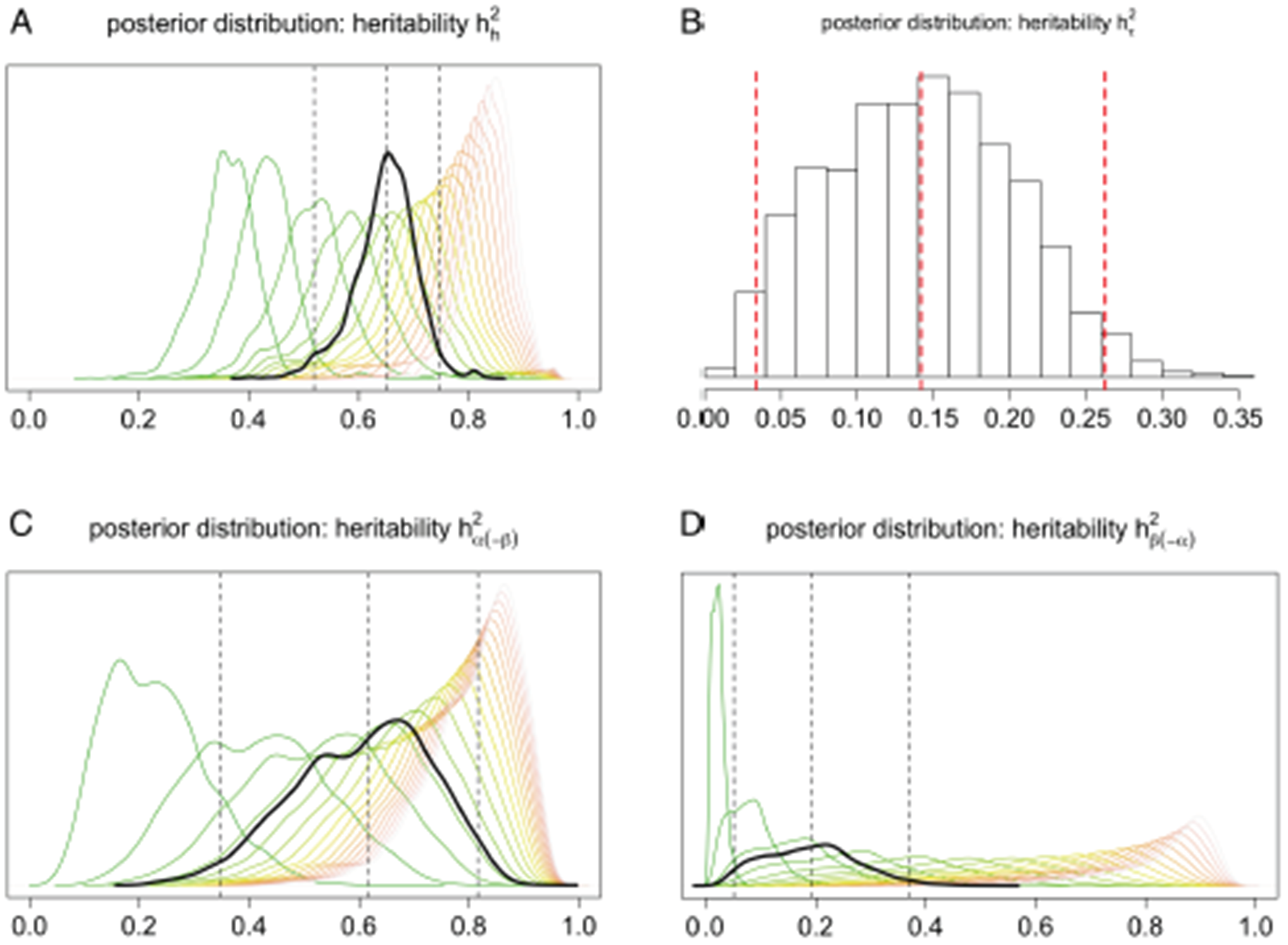
Posterior distributions of pooled measures of heritability (black), pertaining to. (**A**) overall root architecture, (**B**) the near-surface intensity, (**C**) the root bulks location (and size), and (D) the root systems decline of penetration; the middle vertical line marks the posterior median (a Bayesian estimate), and the outer lines delimit the 95% credible interval. In panels A, C, and D, pooling corresponds to integrating depth-dependent heritability over all 18 depths via the harmonic mean of the 18 depth-specific heritability values; the posterior distribution of the unpooled heritability at a given depth is shown in shades of “burnt grass,” where more burnt corresponds to greater depth.

Focusing on the genotypic level (Figure 6 in black; Figure 7, panel A), the Bayesian estimate and 95% credible interval for heritability of the intensity function are, respectively, 0.65 and (0.52, 0.75); for that of the near-surface mean count they are 0.14 and (0.03, 0.26); the bulk parameter, 0.62 and (0.35, 0.82); and the exploration parameter, 0.19 and (0.05, 0.37).

**Figure 7:**
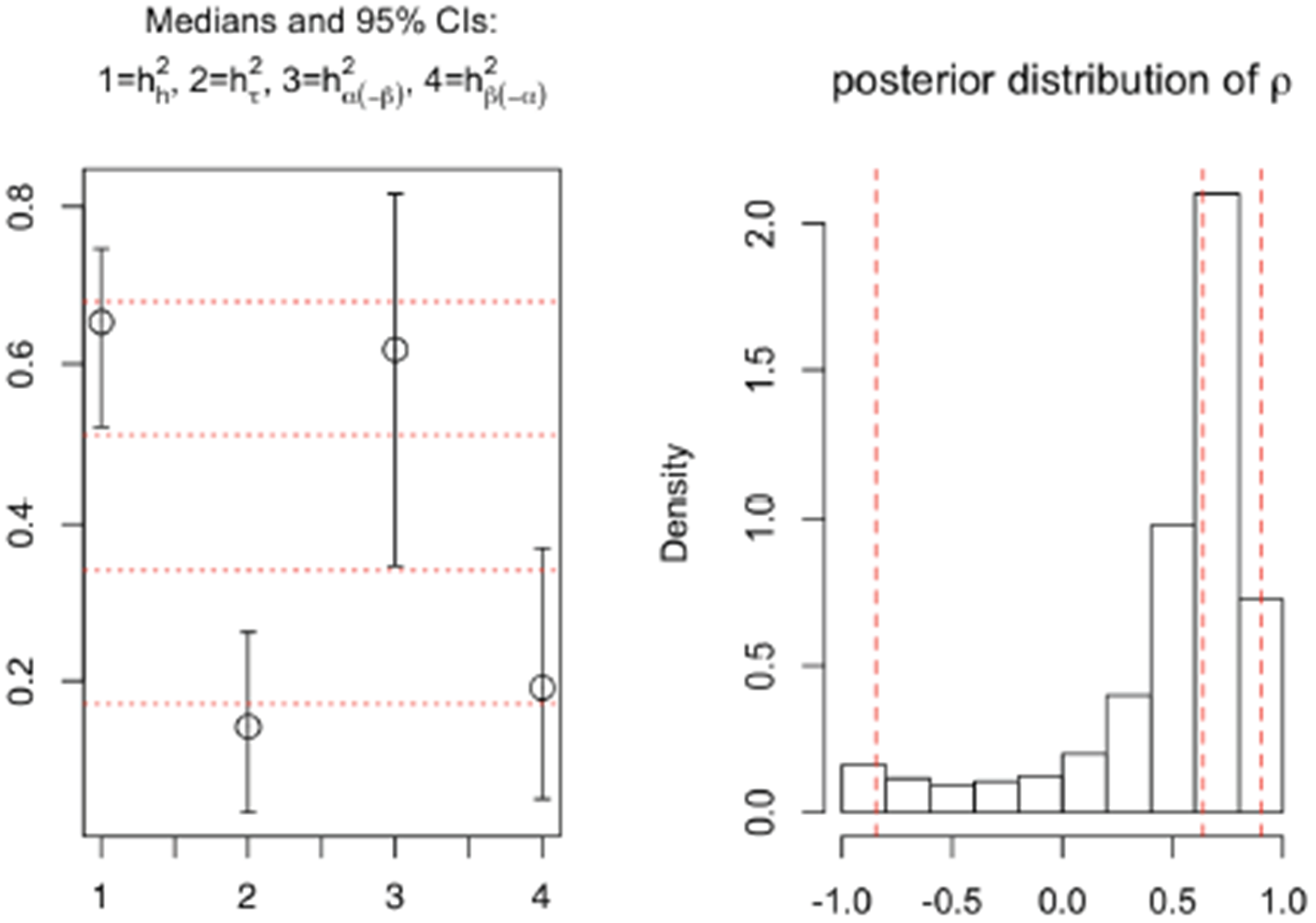
(**A**) 95% credible intervals and posterior medians for the four heritability measures (1=root architecture; 2=near-surface mean count, log scale; 3=bulk parameter; 4=exploration parameter). (**B**) Posterior distribution of *ρ*, with vertical lines indicating the 2.5%, 50%, and 97.5% quantiles.

Note in Figure 6 that (i) the depth-specific components of each of 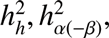 and 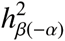 tend to increase as depth increases, and (ii) the near-surface intensity of root count has low heritability (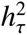). These features of our results indicate that root count features at deeper depths are more heritable than those at shallower depths. In other words, our results provide quantitative rigor for three ideas: the heritability of root architectural traits varies substantively across depth; traits that are associated with a deeper spatial location tend to be more informative about plant genetics; and the depth at which the root system develops its bulk is negatively associated with its tendency to explore deeper. It is also interesting to note that, although overall heritability 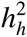 is constituted from 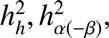 and 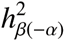, panel A of Figure 7 (which summarizes the genotypic aspects of Figure 6) suggests that each of the latter three tends to be less than 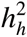 itself, thus root architecture on the whole tends to be more heritable than any of these standalone features of the root system.

#### 3.1.3. Linkage between near-surface root density development and downward exploration

The modeled correlation, *ρ*, between the bulk and exploration parameters (both on the log scale), is estimated to be 0.64, with 95% credible interval = (–0.85, 0.90) (see Figure 7, panel B). Due to skewness, the posterior probability for *ρ* to be positive is 0.88, substantiating that the root systems bulk and downward exploration are generally positively associated architectural features. Specifically, a shallower and more concentrated bulk (small *α*) is associated with a larger tendency for the root system to explore deeper (small *β*). This phenomenon may be regarded as *a small β canceling out a large α*, or, the tendency of exploring downwards to exhibit the effect of negating the tendency to develop root density further away from the surface. We elaborate on this discovery in Section 4.

### 3.2. Validating Model 1

#### 3.2.1. Predictive performance

Although more complex models typically follow the data more closely, they may have poorer predictive performance due to potential overfitting. We consider the predictive performance of *Model 1* by comparing its value of the Watanabe-Akaike information criterion (WAIC) [18] to those for the simpler *Models 2* and *3*, both nested within *Model 1*. As mentioned in Section 2.4 above, the modeled correlation between the bulk and the exploration parameters on the log scale was prespecified as *ρ* = 0 in *Model* 2; *Model 3* considers all model parameters (including those that are genotype-specific) as fixed effects by naïvely prespecifying

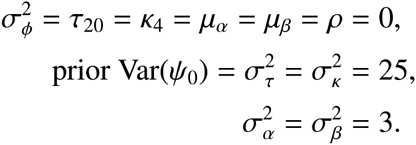

Note that the values 3 and 25 for the prior variance of*ψ*_0_, *τ*_*i*_, *κ*_*j*_, *α*_*i*_, or *β*_*j*_ constitute informative prior distributions for these fixed effects. Before considering 3 or 25, we had specified 10^3^ or 10^4^ for diffuseness. However, in either case, *Model 3* failed to converge due to a weakly identifiable *α*_8_. Consequently, we decided to employ the more restrictive (but defensibly so) prior variances of 3 and 25 according to the following argument.

Based on ref. [25], we deduce that for a wheat plant, the total number of roots at any given depth has a magnitude that is o(100), and thus a generous approximation for the standard deviation (SD) of *θ*_*ijt*_ is 100, or for SD(log *θ*_*ijt*_) is log 100. For *Model 3*, note that

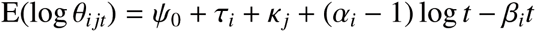

where the sets {*τ*_*i*_}, {*α*_*i*_}, {*β*_*i*_}, and {*k*_*i*_} each follows a linear constraint. Thus, for the priors of the fixed effects, heuristically we let

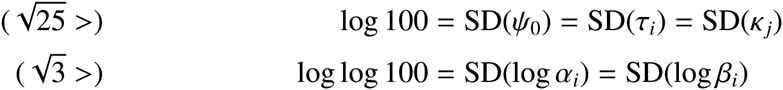

for all *i* ≠ 20 and *j* ≠ 4.

For each *of Models 1-3*, we computed the WAIC based on ref. [26] (see Supplementary Material); they appear in Table 1. There, one can see that predictive performance improved drastically (WAIC decreased from > 46000 to < 35000) from the naïve fixed-effects Model 3 to the mixed-effects Models 1 and 2, both of which are much more complex. Between the two complex models, although Model 1 is slightly larger than Model 2 (by a single parameter that represents a priori dependence between the bulk and exploration parameters), effectively they perform equally well in predictive power, as suggested by a merely nominal difference (=3) in WAIC values.

**Table 1:** Values of the Watanabe-Akaike information criterion (WAIC) as a measure of predictive performance by our Bayesian HNLMMs. A smaller WAIC value suggests better model performance.

| Model | Description | WAIC |
| --- | --- | --- |
| 1 | Most sophisticated among our models, as presented in this paper | 34198 |
| 2 | Same as *Model 1*, but with a prespecified $\rho = 0$ | 34195 |
| 3 | Naïve preliminary model: same as *Model 2*, except with $\phi_{ijt}$ term missing and all of $\psi_0, \tau_i, \alpha_i, \beta_i$, and $\kappa_j$ taken as fixed effects | 46627 |

In addition to a mere nominal difference in WAIC values between Models 1 and 2, we also observed that the approximate values for the effective number of parameters, p_WAIC_ (used in the computation of WAIC), increased from 4011 for Model 1 to 4014 for Model 2. Importantly, although model complexity was increased from Model 2 to Model 1 by correlating the genotypic bulk and exploration parameters through a study-wide parameter, the additional parameter reduced the models overall amount of unknownness. Thus, these values of *p*_WAIC_, along with those of the WAIC and the substantively large posterior probability that *ρ* > 0, suggest the merit of retaining ρ as an unknown parameter in the hierarchical nonlinear mixed model (HNLMM).

In summary, disregarding the hierarchical structure between the root count intensity function and its various random components that are specific to genotypes, plots, and depths led to a naïve model that agreed poorly with the empirical behavior of our root count data. For the hierarchical models, predictive performance remains effectively unaffected whether a priori dependence between the bulk and exploration parameters is considered; however, we regard this extra dependency as a key biological feature because it improves the interpretability of the model by providing an explicit assessment of the interplay between the bulk and exploration parameters, and this in terplay is shown to be substantive based on our field data.

#### 3.2.2. Residual Plots

Next, we inspect violin plots produced by the R package ‘vioplot’ [27] for residuals that correspond to the Level 1 noise terms *τ*_*i*_ and *φ*_*ijt*_ in *Model 1*. Non-noise-like patterns in these residual plots would suggest the statistical inadequacy *of Model 1* for our data.

Figure 8 is based on the posterior distribution of *φ*_*ijt*_ (posterior median shown as black dot), plotted against the posterior mean of log *θ*_*ijt*_ rather than the log-transformed *ȳ*_*ij+t*_ = Σ_*k*_ *y*_*ijkt*_/4. This is because *ȳ*_*ij+t*_ = 0 for 181 out of all 1440 combinations of {*i,j,t*} (see Section 4 for possible implications). Figure 8 shows that *φ*_*ijt*_ has a (a) slight tendency to increase with log*θ*_*ijt*_, and (b) a distinctive non-random relationship with log*θ*_*ijt*_ especially when the latter is small (which is typically at lower depths). We break down this relationship by the residual violin plots in **Figures 9-13**.

**Figure 8:**
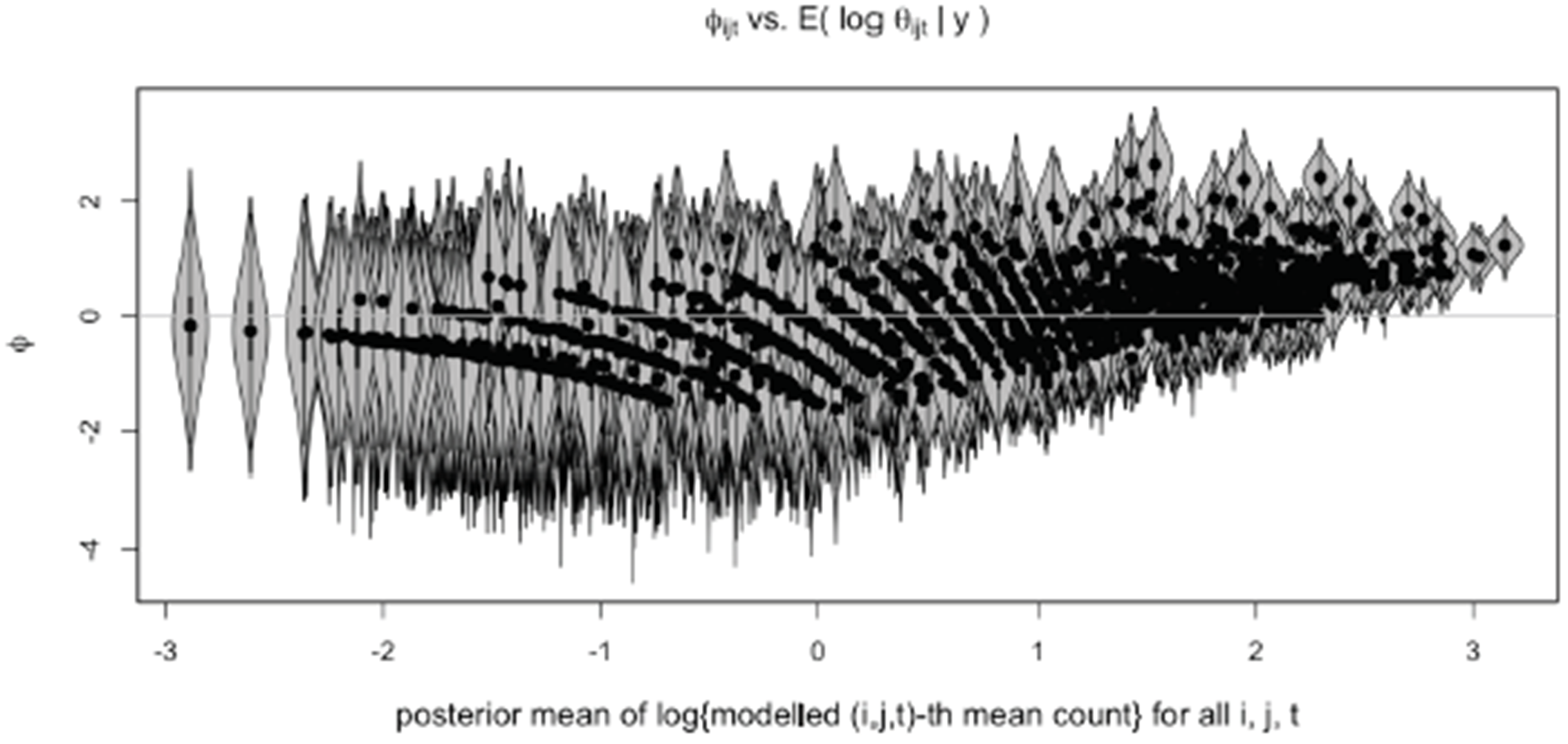
Violin plots: Level 1 noise *Ф* against the posterior mean of log(modeled mean root count), for all {i, j, t} combinations. The median that corresponds to each {i, j, t}-th violin of *Ф* is shown in black. Some non-random patterns are noticeable.

Overall, we observe the following minor anomalies:

- *τ*_*i*_ vs. *ȳ*_*i++t*_ (Figure 10): at many depths *t*, the random effect *τ*_*i*_ has a slight tendency to be negative for small observed values of *ȳ*_*i++t*_, and positive for large observed values of *ȳ*_*i++t*_;
- *Ф*_*ijt*_ vs. *i* (Figure 11): at *t* = 160,170, or 180 cm, the residual *Ф*_*ijt*_ for a small number of plots ({*i, j*} combinations) has a tendency to be highly positive;

**Figure 10:**
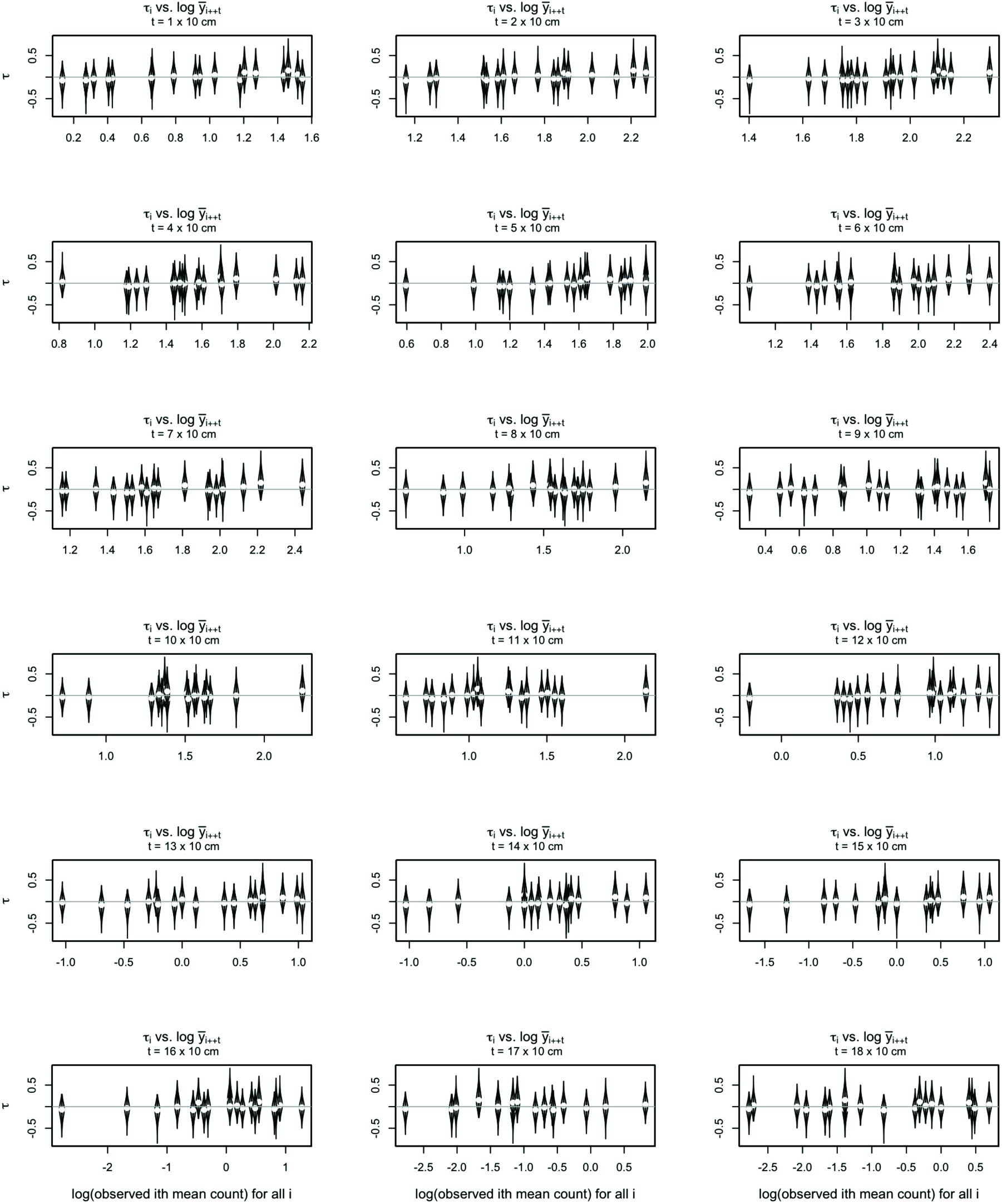
Violin plots: random effect *τ* against log(mean count); each panel is associated with a particular depth *t*. Posterior medians of *τ* are shown in white. A slight increasing trend is noticeable at various values of *t*.
- *Ф*_*ijt*_ vs. *t* (Figure 12): for the 3rd replicate block (*j* = 3), many genotypes (e.g. *i* = 2,12,14, etc.) at the deepest six depths are associated with (*Ф*_*ijt*_ that increases with depth systematically; the same applies to *j* = 4 and *i* = 8,9,11, etc.; additionally, the plot {*i, j*} = {5,3} shows that *Ф*_*ijt*_ has a tendency to be all positive;
- *Ф*_*ijt*_ vs. *j* (Figure 13): the same conclusion as for Figure 11 for *t* = 160,170, or 180 cm; additionally, *Ф*_*ijt*_ for a small number of plots has a slight tendency to be positive at *t* = 30 or 100 cm.

**Figure 11:**
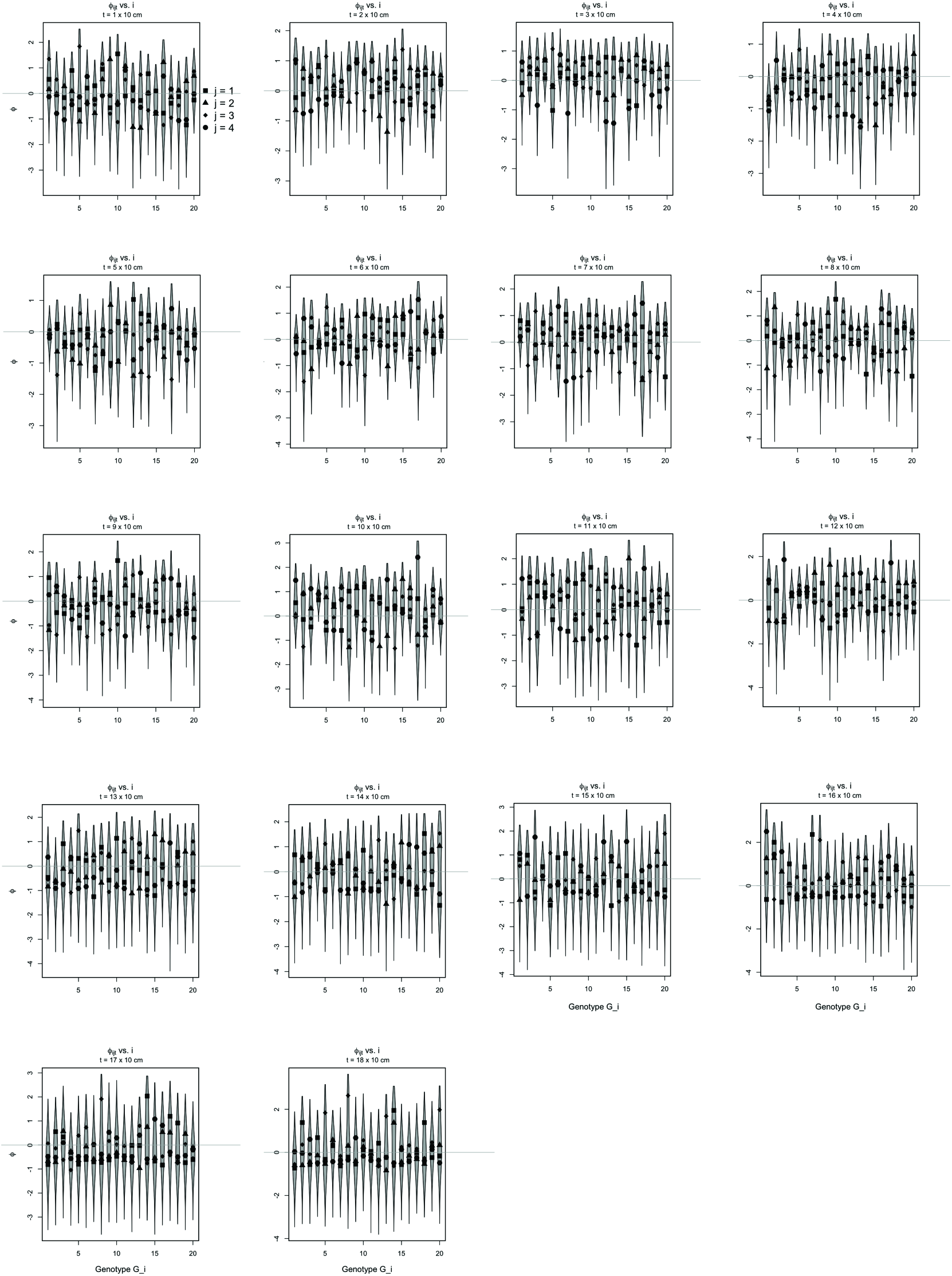
Violin plots: Level 1 noise *φ* against genotype i; each panel is associated with a particular depth *t*. Inside the tth panel are 20 genotype-specific violin plots, each {i, t}-th violin corresponding to pooling four plot-specific violins (over blocks *j* = 1, 2, 3, 4). The plot-specific medians of *φ* are shown in black (see top-left panel for legend). Minor anomalies are apparent at the highest values of *t*.

**Figure 12:**
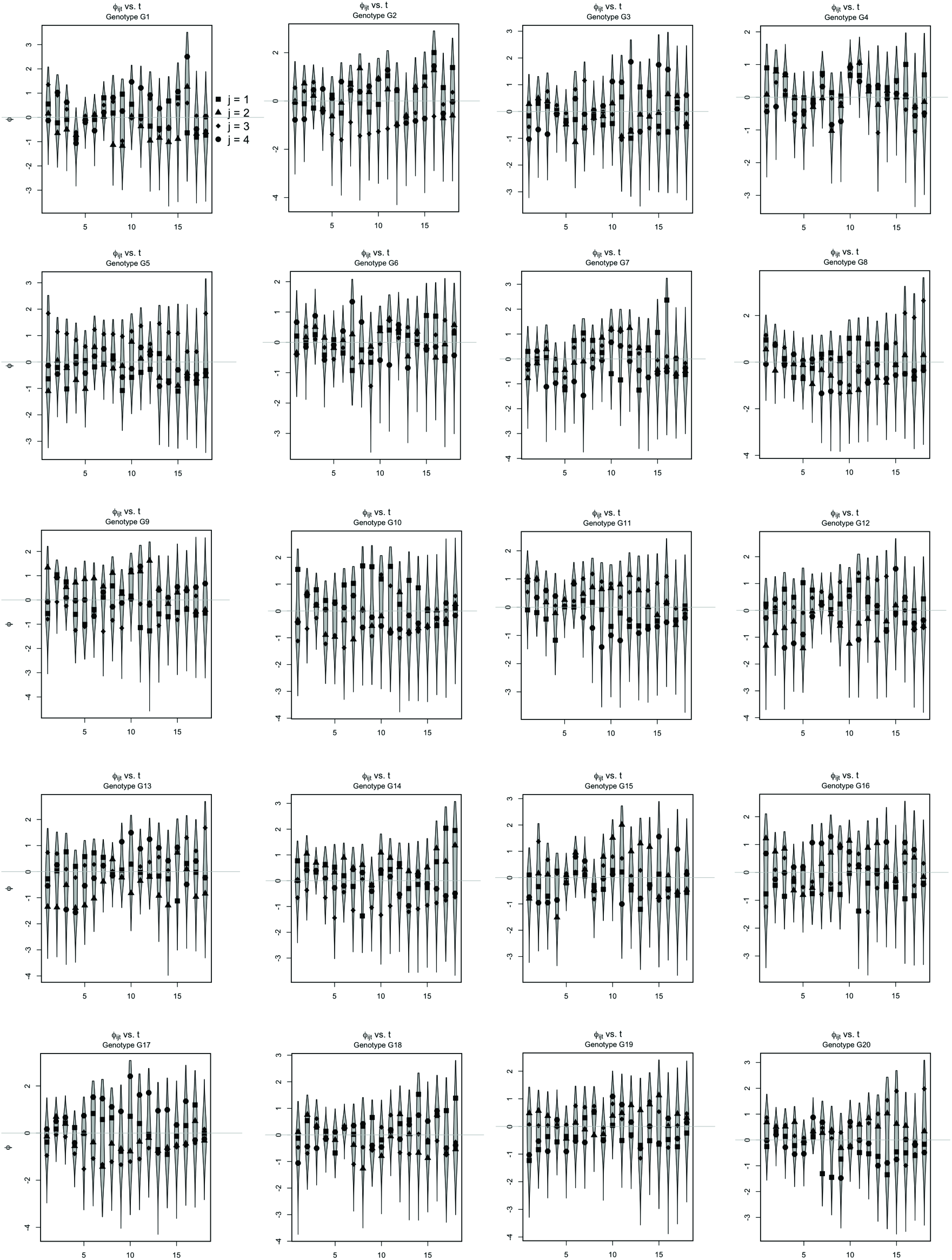
Same as Figure 11, except for the role reversal between genotype *i* and depth index *t*. Minor anomalies over higher values of *t* are apparent at isolated values of *i* and/or *j*.

**Figure 13:**
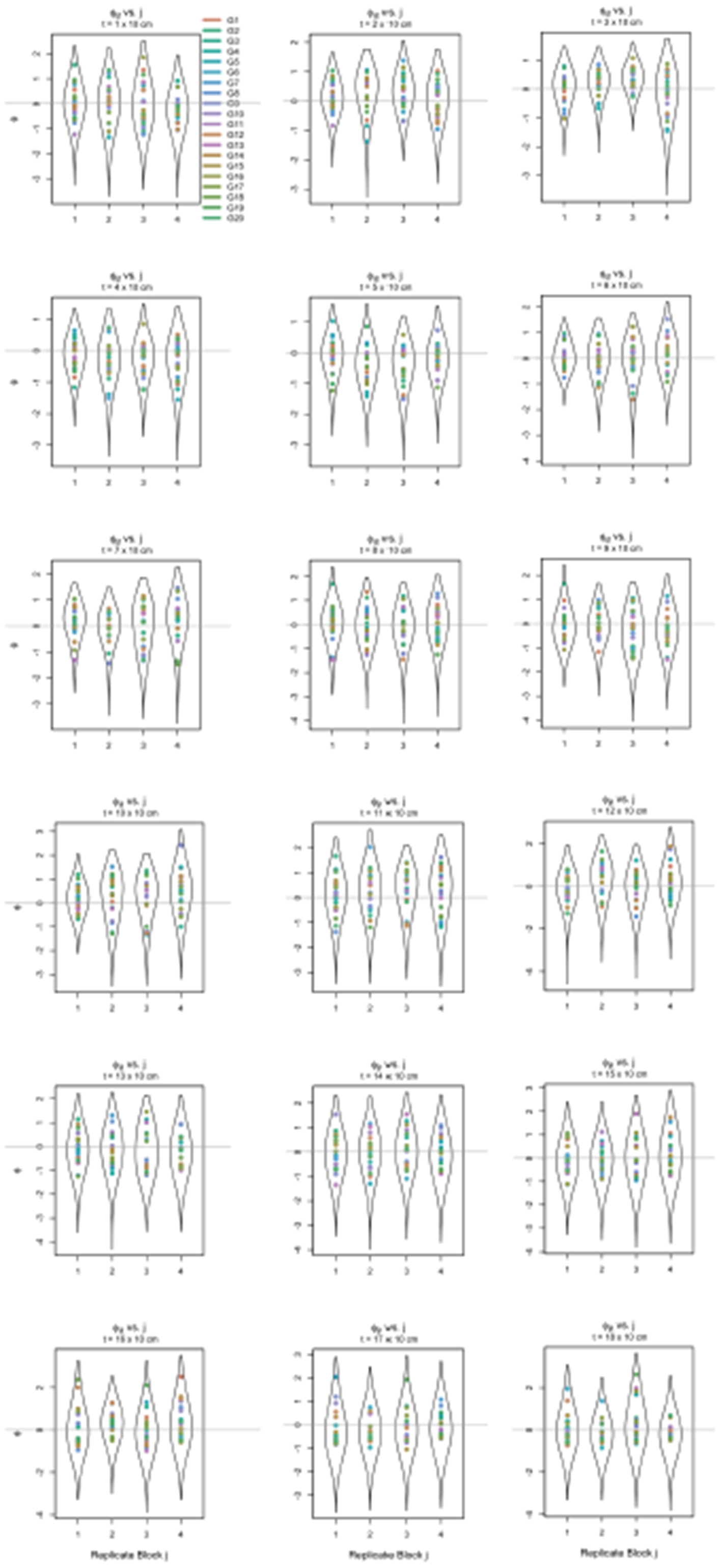
Violin plots: Level 1 noise *φ* against replicate block *j*; each panel is associated with a particular depth *t*. Inside the tth panel are 4 block-specific violin plots, each {*j, t*}-th violin corresponding to pooling 20 genotype-specific violins (over genotypes *i* = *G*1, …, *G*20). The genotype-specific medians of *φ* are shown in color (see top-left panel for legend). Minor anomalies are apparent at various values of *t*.

Altogether, the residual violin plots suggest that the statistical inadequacy of *Model 1* lies in the modeled behavior of *θ*_*ijt*_ across the deeper depths for specific combinations of {*i, j*}. In Section 4 we provide an overview of possible directions that may be taken to improve the adequacy of our HNLMM.

**Figure 9:**
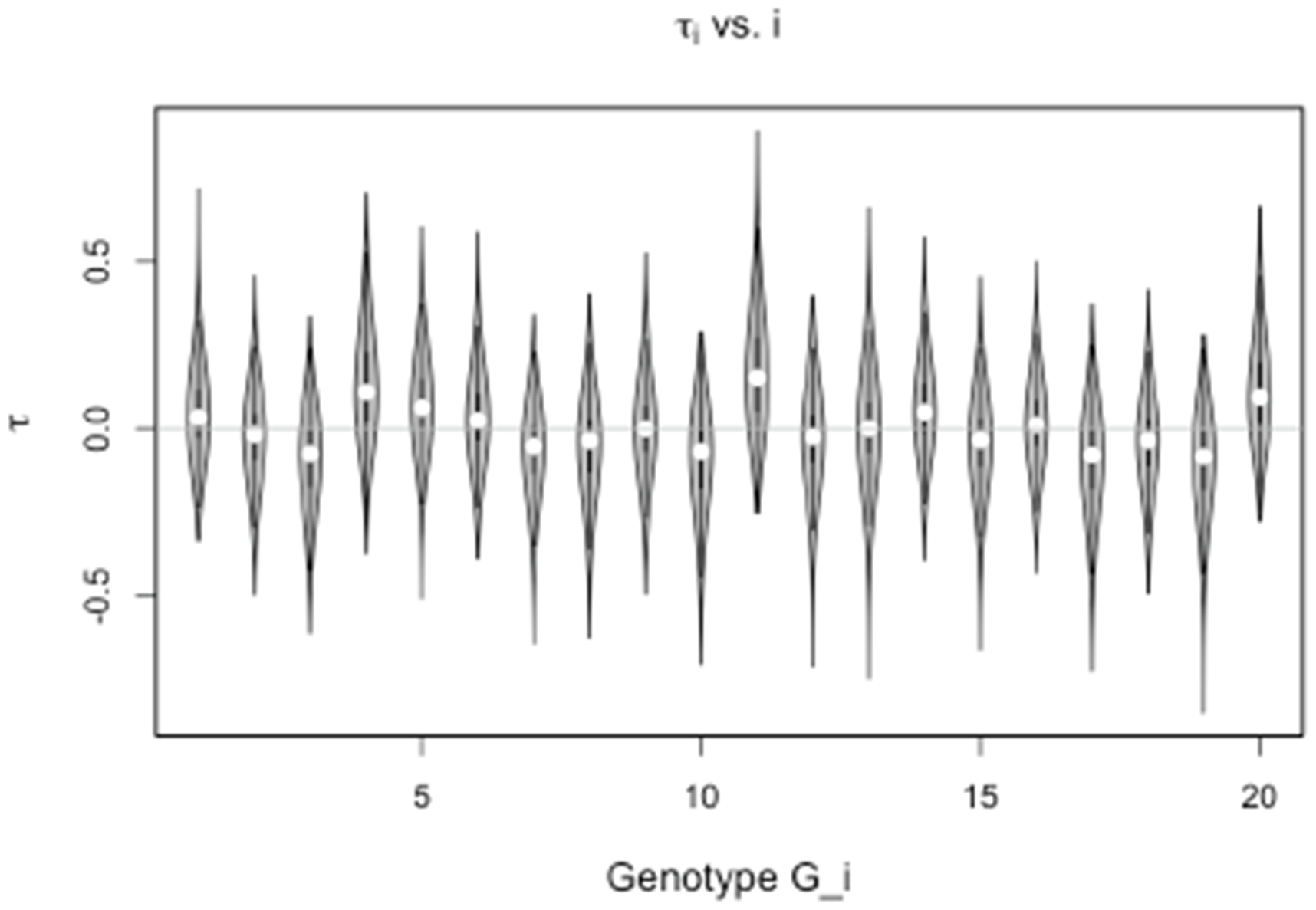
Violin plots: genotypic random effect *τ* against genotype *i*. Each ith violin also shows the posterior median of *τ* in white. No anomalies are apparent.

## 4. Discussion

The development of roots in response to the extreme heterogeneity of the soil results in a lack of discernible root system characteristics that can be measured in the field and integrated into crop breeding programs. In this paper, we have addressed this lack by scrutinizing root counts observed using a core-break count method, and by developing a novel modeling approach that accounts for all root count data holistically. Our approach gave rise to new multiresolution heritability metrics, each describing a specific feature of the root count distribution spatially and at the overall genotypic level, which we showed to be substantially heritable. Our integrative approach can allow selective pre-breeding programs for root distribution and may facilitate the identification of genetic markers from field data.

The holistic nature of our approach is an intrinsic advantage of Bayesian hierarchical modeling. Sufficient computational resources and algorithms can often easily handle Bayesian inference from models and data structure of arbitrary complexity. In contrast, classical statistical inference can be too impractical when models or data structures deviate from well-studied scenarios. In our case, the experimental setup and the notion of root architecture together led to a highly non-standard scenario that, under a classical paradigm, would have been much less straightforward to model and subsequently draw inference from. Not only was *y* (the response variable of interest) strictly non-Gaussian even under any reasonable variable transformation, the data were also three-dimensionally spatial in nature, where replicate plots were arranged in a certain 2-dimensional structure (indexed by {*i, j*}), and in turn each plot generating numerous 1-dimensional spatial observations of *y* (indexed by *t*).

With Bayesian inference, model diagnosis/validation is also straightforward to conduct. The satisfactory predictive power of our Model 1, as described earlier, suggests that Model 1 is scientifically sensible and has yielded biological insights that are superior to what could have been drawn from previous linear models applied to core-level metrics (from collapsing segment-level data).

Although Model 1 does not account for potential within-core spatial dependency among segment-level root count data (see Section 4.1 below), the biological implications of this model nonetheless will help to define root traits for breeding. The canonical model of root distribution with depth is that of a negative exponential function [28]. Ref. [29] describes a nonlinear function Y = 1 - *β*^d^ where Y is the cumulative root fraction from the surface to the depth (*d* cm), and the coefficient *β* is genotypically determined. This model was later employed in ref. [30] to model root distributions across a range of terrestrial biomes. However, this 1-dimensional model takes no account of the horizontal distance from the base of the plant. It has been observed that root distribution is 1-dimensional with depth in grassland, 2-dimensional in crops planted densely in rows, and 3-dimensional where plants are widely spaced [31]. The simulation studies in ref. [32] showed that root length density (RLD) — typically the length of root per volume of soil (cm/cm^3^) — for maize decreased nonlinearly with horizontal distance from the stem in the top 40 cm, but below that depth they were homogeneously distributed with horizontal distribution from the plant. Ref. [32] also showed that the roots were subject to clustering at all depths, and that whilst there was no preferential growth in a horizontal plane, the orientation of root growth deviated from the uniform distribution with increasing depth. Similar findings were generated in the simulation study in ref. [33], and both studies drew attention to the likely effect of soil structure to further perturb the uniform directional distribution of root development parameters.

The similarity between the model in ref. [29] and the special case of our gamma kernel function where *α* = 1 should be noted (Figure 3, panel A). Rather than RLD, our model accounts for root counts that are random with respect to sampling position by row in a crop. The model is designed to explain the distribution of root counts with depth at the crop level (and not the plant level). However, the sampling position is likely to have a strong influence on the surface root counts, which explains the low heritability of *τ* in our model.

Interpreting the biological meaning of the *bulk* and *exploration* parameters (*α* and *β*, respectively) is also interesting. In the gamma kernel function, *β* also affects the depth and intensity of the peak otherwise defined by *α* (Figure 3). Indeed, our data analysis implicated that *α* and *β* were positively correlated (Figure 7, panel B). For our HNLMMs, predictive performance remained largely unaffected whether *a priori* dependence between *α* and *β* is considered; however, including this extra dependency improved the interpretability of the model by providing an explicit assessment of the interplay between the root systems tendencies to branch beneath the surface and to explore vertically, deep below the surface.

An explanation for this effect may be found in the structure of the soil; root growth in deeper layers is perceivably constrained to networks of cracks and pores [34]. Ref. [16] shows that in a dense, structured subsoil 85-100% of roots below 60 cm were clumped in pores and cracks in the soil (compared to 30-40% above 60 cm), and 44% of the roots were clumped in pores with more than three other roots. Exploration of the soil for cracks and pores may define the exploitation of the soil by a root system. It has been suggested that plants have evolved randomness and instability in their root system development [35], which may facilitate exploration. The exploration of the shallow layers for cracks and pores may be what determines the eventual depth; our model implies that more branching near the surface gives better access to the subsoil.

The primary purpose of our Bayesian model was to distinguish genotypes from root count data that are statistically noisy. The inference for heritability based on the intensity functions suggests that our approach can be used to identify genetic markers of root system distribution in field data; identified markers then could be integrated into breeding programs. The high heritability of the “bulk” parameter also suggests that a breeding program could successfully alter the depth at which a root system proliferates.

Notwithstanding, residual plots suggest some minor statistical inadequacies of *Model 1*. Therefore, it may be advantageous to (1) formally model the within-core spatial dependence (possibly at a higher spatial resolution of core depths than the current 10 cm intervals) and (2) also incorporate an additional two-dimensional spatial correlation structure among field plots. See Section 4.1 below for practical implications of modeling such 3-dimensional spatial dependence.

Finally, it may also be of benefit to develop a new quantitative framework to predict RLD from the posterior mean root count profiles while accounting for trials in different soil and climate conditions, under which the response of the intensity functions and their underlying parameters to subsoil constraints could be rigorously exploited.

### 4.1. Technical remarks

Note that while 5 cm segments were produced in the field, the first, third, fifth, etc. depths were ignored in the statistical modeling; only imaged counts at depths in 10 cm increments from the surface were considered. This implies that within the same core, the resulting counts *y* were less spatially autocorrelated due to a lower spatial resolution of the data from omitting alternate segments from consideration. With segment-level counts *y* thus produced, we considered for statistical modeling 18 (= *n*_*D*_) depth values from each core, from depth 10 cm to depth 180 cm.

Also note that our mathematical expression of *Model 1* and its variants employs a convention for time series analyses in which depth *t* (an analog of time) is effectively an integer index ranging over 1 to 18. In practice, an alternative parametrization may be preferred, such as *t* = 10, 20, …, 180 cm, or even to mathematically map the observed spatial domain to the unitless interval (0, 1] so that the labels are *t* = 1/*T*, 2/*T*,…, (*T* − 1)/*T*, 1 where *T* is the maximum number of core segments. We refer to the latter as the canonical scale for depth, for which we discuss as follows the invariance of our model inference whether the depth scale employed in practice is canonical or otherwise.

In general, as long as the root counts are observed at regular spatial intervals along a soil core, the set of labels

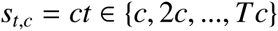

is possible in practice for some *c* ≠ 0. However, does different values of *c* result in different statistical inference?

The answer is “no.” Note that the intensity functions kernel is

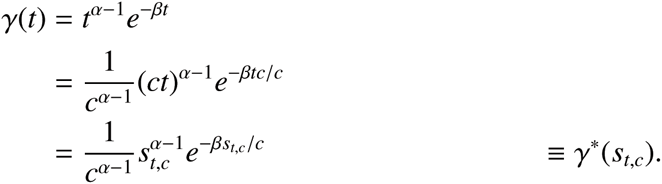

We define the *canonical scale* for depth as *s*_*t,c*_=1/*T*, so that

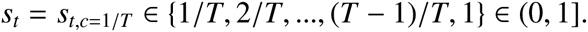

Thus, on the canonical scale, the formulation *of Model 1* remains the same except for

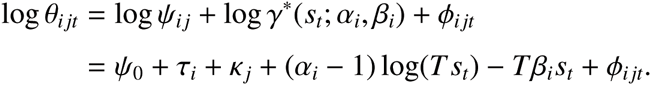

This reparametrization using *γ\** is linear in both *s*_*t*_ and *T* = 1/*c*, and thus the statistical inference is invariant to any of the conventional depth scale *t*, the reparametrized scale involving a non-zero *c*, or the canonical scale *s*_*t*_. This invariance is similar to that in ref. [36] where the rigor of the statistical inference is developed on the canonical scale rather than the conventional scale.

Finally, we discuss possible extensions to *Model 1*. Although the hierarchical structure of our HNLMM already addresses some over-or under-dispersion in the raw root counts, an abundance (181/1440 > 12%) of 0 mean root counts (averaged over four replicate cores) suggests a potential need to include a formal zero-inflation component in a future improved model (e.g. via a mixture model). Furthermore, at present our HNLMM does not include formal spatial statistical modeling [37] of within-core spatial dependency among segment-level root count data at a 10 cm spatial resolution. Spatial statistical modeling would impose substantial complexity to the statistical inference and computational burden. However, the residual plots perhaps suggest that formal spatial modeling could be a valuable additional component for our HNLMM, especially with data at the 5 cm resolution. A possible spatial structure could be an autoregressive dependence over depth, and/or a nearest-neighbor dependence (among field plots) that constitutes a Markov random field [37].

## Acknowledgments

We thank Prof. M. Watt (Institute of Bio-and Geosciences 2, Forschungszentrum Jülich) for support and guidance. We also thank Bayer CropScience for (a) funding CSIRO for this research project, part of which was subcontracted to the Australian National University to sponsor G.S.C.’s involvement after June 2015; and (b) sponsoring the CSIRO Agriculture Vacation Scholarship 2014 that was held by T.R.B., under which he contributed to the basic model that was later extended to the approach in this paper. G.S.C. thanks the University of Washington and University of Waterloo for online resources made available to her as an affiliate/adjunct faculty member.

## Supplementary Material

Our dataset and computer code will be made available through the journal upon acceptance of this manuscript for publication.

